# In vivo Restricted-Diffusion Imaging (RDI) is sensitive to differences in axonal density in typical children and adults

**DOI:** 10.1101/2020.08.17.254656

**Authors:** Dea Garic, Fang-Cheng Yeh, Paulo Graziano, Anthony Steven Dick

## Abstract

**Background:** The ability to dissociate axonal density *in vivo* from other microstructural properties of white matter is important for the diagnosis and treatment of neurologic disease, and new methods to do so are being developed. We investigated one such method–restricted diffusion imaging (RDI)–to see whether it can more accurately replicate histological axonal density patterns in the corpus callosum (CC) of adults and children compared to diffusion tensor imaging (DTI), neurite orientation dispersion and density imaging (NODDI), and generalized q-sampling imaging (GQI) methods. To do so, we compared known axonal density patterns defined by histology to to diffusion-weighted imaging (DWI) scans of 840 healthy 20- to 40-year-old adults, and, in a replication and extension, to DWI scans of 129 typically developing 7-month-old to 18-year-old children and adolescents. Contrast analyses were used to to compare pattern similarities between the *in-vivo* metric and previously-published histological density models. We found that RDI was effective at mapping axonal density of small (Cohen’s *d*= 2.60) and large fiber sizes (Cohen’s *d*= 2.84) in adults. The same pattern was observed in the developing sample (Cohen’s *d*= 3.09 and 3.78, respectively). Other metrics, notably NODDI’s intracellular volume fraction (ICVF), were also sensitive to differences in axonal density across the longitudinal axis of the CC. In conclusion, the study showed that RDI is effective at measuring axonal density of small and large axons in adults and children, with both single- and multi-shell acquisition DWI data. Its effectiveness and availability to be used on standard as well as advanced DWI acquisitions makes it a promising method in clinical settings.

## Introduction

The ability to quantify microstructural properties of neural tissue is important for the diagnosis and treatment of neurologic disease. Significant advances in the acquisition and analysis of diffusion weighted magnetic resonance imaging (DW-MRI) data have allowed researchers to target specific sources of the DW-MRI signal associated with specific tissue properties. Recent analysis approaches, such as restricted diffusion imaging^1^ and neurite orientation dispersion and density imaging^2^ have provided more fine-grained metrics that augment, and in some cases replace, the more traditional metrics that are reported in diffusion tensor imaging (DTI) studies. The present investigation is an attempt to understand whether such new metrics are sensitive to a particular microstructural property of white matter, that of axonal density. Providing an *in vivo* measure of axonal density has high potential for clinical importance. For example, histological studies have shown that axonal density is related to the progression rate of multiple sclerosis^3^, hereditary spastic paraplegia^4^, and it is a sensitive measure of multiple sclerosis lesion in an animal model^5^. The density of fibers of a particular diameter is also important because neural conduction velocity scales with axonal diameter^6^, influencing the timing and rapidity of information transfer in the nervous system. Thus, being able to quantify these density changes *in vivo* during development could provide essential insight into disease progression, development, and general nervous system structure/function relationships.

The new metrics we investigate extend from previous algorithms used to reconstruct DWI data. The most popular instantiation of DWI reconstruction is diffusion tensor imaging (DTI), which obtains familiar metrics such as fractional anisotropy (FA), mean diffusivity (MD), axial diffusivity (AD), and radial diffusivity (RD), that provide information about the directional nature of water diffusion through tissue. AD measures the longitudinal component or eigenvalue (*λ*_1_) of the diffusion tensor model, and it has been shown to be sensitive to axon integrity^7^. RD is the average of the remaining eigenvalues ([*λ*_2_+ *λ*_3_]/2), which are perpendicular to the longitudinal component, and which have been associated with myelin integrity^7, 8^. MD is the average of these three principal eigenvalues ([*λ*_1_ + *λ*_2_ + *λ*_3_]/3). Finally, FA is the most frequently used metric. This metric indicates how the other diffusion metrics stand in relation to one another, providing a summary index of the directional nature of water diffusion in the voxel^9^. It is scaled such that 0 represents unrestricted or isotropic diffusion, and 1 indicates water diffusion that is anisotropic, or restricted in all but one direction. These metrics have been useful in tracking various fiber pathways in health and disease across the lifespan^10–12^, but none of these DTI metrics are designed to isolate and characterize the specific contributions of axonal density to the diffusion-weighted imaging signal.

With updates in MRI hardware and software, more diffusion directions can be acquired in a shorter amount of time, which has allowed for the use of better imaging reconstruction algorithms. These High Angular Resolution Diffusion Imaging (HARDI) algorithms improve the estimation of water diffusion in multiple directions, leading to better estimation and resolution of such crossing and kissing fibers. For example, Generalized Q-Sampling Imaging (GQI) has been employed to efficiently reconstruct the orientation distribution function (ODF) of water diffusion from HARDI acquisitions^13, 14^. Two metrics can be recovered from these diffusion models: Quantitative anisotropy (QA) and generalized fractional anisotropy (GFA). QA is defined as the amount of anisotropic spins that diffuse along a fiber orientation, while GFA can be thought of as a higher-order generalization of FA^15, 16^. Regardless of the improvements GQI has provided for MRI, these metrics are still not optimal for measuring axonal density differences^2^.

Other imaging reconstruction methods have focused on estimation of microstructural properties of cell body processes—dendrites and axons, i.e., neurites. For example, Neurite Orientation Dispersion and Density Imaging (NODDI) ascribes the signal of water protons in tissue into three components: isotropic diffusion representing free motion in areas such as ventricles, intracellular volume diffusion representing restricted water in dendrites and neurons, and extracellular volume diffusion representing glial cells, cell bodies, and the extracellular environment^2^. A number of studies have compared the NODDI signal to known histological patterns in white matter. For example, Zhang et al^2^ have shown that NODDI is capable of disentangling the two components of FA, neurite density and orientation dispersion, allowing the two to be studied individually. In an important study that serves as a model for our present investigation, Genc et al^17^ have also used NODDI to map corpus callosum density in children and adolescents. They found that NODDI intracellular volume fraction (ICVF) and orientation distribution function (OD) metrics were sensitive to histologic differences along the longitudinal axis of the corpus callosum. We conducted a partial replication of this study, and will return to the details of it in the Method.

Finally, Restricted Diffusion Imaging (RDI) is a recent measure that has been proposed to index changes in cellularity (i.e., cellular density), which affects local diffusion of water molecules. RDI attempts to separate non-restricted diffusion of water molecules from restricted diffusion of water molecules by focusing on the difference in diffusion displacement of each component. As such, the model can measure restricted diffusion while ignoring non-restricted diffusion. Yeh et al.,^1^ showed that RDI was sensitive to differences in cellular density in a manufactured phantom, and in rat cardiac tissue that sustained a lesion, which resulted in the migration of macrophages to the lesion site. The density changes resulting from this macrophage migration were detectable in changes in RDI. Thus, RDI seems to be sensitive to cellular density and changes in cellular density.

What is unknown, though, from these initial studies is whether RDI metrics are sensitive to density of axonal fibers. It is well known that axons at cross-section differ in diameter^18^ and this affects the density of axons within the fiber pathway. For example, the Aboitiz et al.^18^ histological examination of 20 post-mortem healthy adult brains showed that the density of small and large diameter axons differs across the anterior-posterior axis of the corpus callosum. This now well-established density pattern has been replicated with other measures in more recent work^19, 20, 21^, including the study by Genc and colleagues^22^ that serves as a model of the present investigation. As we noted earlier, it would be important for a number of reasons to establish *in vivo* metrics that are sensitive to differences in axonal density. Given its high sensitivity to diffusion changes as a function of cellular density, we expected RDI to be most sensitive to these changes relative to other established metrics. Thus, the primary aim of this study is to address whether RDI is a candidate metric for measuring axonal density in vivo, and further whether it does so better than other available diffusion metrics. To do this, we measured segments of the corpus callosum, which have been known to differ in axonal density on the anterior-posterior axis^18, 19, 23, 24^. We used both a healthy adult sample (n = 840) and a healthy developing sample (n = 129) to test whether previous histological results match the segmental pattern reflected using RDI in vivo. In order to demonstrate that RDI measures a unique characteristic in the imaging modality, we compared the RDI measurements to that of DTI metrics (AD, RD, FA, and MD), GQI diffusion metrics (GFA and QA), and NODDI metrics (ISO, ICVF, and OD). We hypothesized that RDI would most reliably, relative to other measures, replicate the Aboitiz^18^ anterior-posterior density pattern established in histology.

### Paradigm and Summary of Both Experiments

Two studies were designed to investigate the sensitivity of various diffusion metrics to the anterior-posterior microstructural organization of the corpus callosum. Our study design uses as a point of departure the study by Genc et al^22^. We used two publicly available datasets. In Study 1 we analyzed 840 multi-shell adult high angular resolution diffusion-weighted imaging (HARDI) scans from the Human Connectome Project (http://www.humanconnectome.org). Study 1 was designed to replicate and match the corpus callosum density pattern found in the Aboitiz et al^18^ histological study. In Study 2, we conducted the same analysis on a different dataset of 129 infants, children, and adolescents from the C-MIND study (https://research.cchmc.org/c-mind/). We used the single-shell acquisition from this study to establish whether the RDI metric could be sensitive in single-shell data, which are also still common acquisitions, especially in clinical settings. Simultaneously, the use of the CMIND data allowed us to investigate whether the density pattern can be detected in a developmental sample. For both studies we computed DTI metrics (FA, RD, AD, MD), GQI metrics (QA, GFA, and RDI), and (in Study 1 only) NODDI metrics (ICVF, OD, ISO). We applied a corpus callosum mask based on Aboitiz et al.^18^ to compute summary statistics of each metric for each subregion. The mask was applied in an automated fashion for Study 1, and hand-drawn for Study 2 (to accommodate the different brain sizes in the developmental sample^25^).

## Study 1

### Participants

The adult sample contained 842 participants between the ages of 22- and 35-years-old who underwent MRI scans as part of the Human Connectome Project. Data were provided by the Human Connectome Project, WU-Minn Consortium (Principal Investigators: David Van Essen and Kamil Ugurbil; 1U54MH091657) funded by the 16 NIH Institutes and Centers that support the NIH Blueprint for Neuroscience Research; and by the McDonnell Center for Systems Neuroscience at Washington University. The automated atlas segmentation was successfully applied to 840 of these participants. Therefore, the summary statistics for the DTI and GQI metrics were calculated on 840 participants. The NODDI reconstruction is a time and resource intensive process, and thus a random sample of 100 participants were selected from the full sample for this reconstruction.

#### Diffusion Metrics

We calculated and compared four DTI metrics (FA, RD, AD, and MD), three GQI metrics (nQA, GFA, and RDI), and three NODDI metrics (ICVF, OD, ISO). The diffusion tensor was used to calculate the eigenvalues reflecting diffusion parallel and perpendicular to each of the fibers along three axes (x, y, z). The resulting eigenvalues were then used to compute indices of FA, RD, AD, and MD^26, 27^. FA is an index for the amount of diffusion asymmetry within a voxel, normalized to take values from zero (isotropic diffusion) to one (anisotropic diffusion). FA is sensitive to microstructural changes in white matter, with higher FA values indicating more directional diffusion of water. This value can be decomposed into AD, measuring the parallel eigenvalue (*λ*_1_), and RD, measuring the average of the secondary and tertiary perpendicular eigenvalues ([(*λ*_2_ + *λ*_3_)]/2). AD and RD quantifications are sensitive to axon integrity and myelin integrity, respectively (Winston, 2012). MD is a summary mean of the three principal eigenvalues ([*λ*_1_ + *λ*_2_ + *λ*_3_]/3). Figure 1 displays the diffusion tensor model under isotropic and anisotropic diffusion.

**Figure 1.**
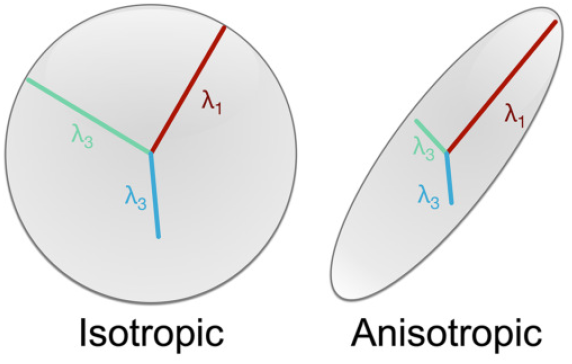
The diffusion tensor model is illustrated/ Left illustrates isotropic diffusion. Right illustrates anisotropic diffusion. *λ* s represent the principal eigenvectors. *λ*_1_ = Longitudinal (axial) diffusivity; [*λ*_2_ + *λ*_3_]/2 = Radial diffusivity. [*λ*_1_ + *λ*_2_ + *λ*_3_]/3) = Mean diffusivity.

Three GQI metrics were also calculated—QA, GFA, and RDI.

##### Quantitative Anisotropy (QA)

QA is defined as the amount of anisotropic spins that diffuse along a fiber orientation, and it is given mathematically by:

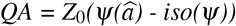

where *ψ* is the spin distribution function (SDF) estimated using the generalized q-sampling imaging, 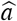 is the orientation of the fiber of interest, and *iso(ψ)* is the isotropic background diffusion of the SDF. *Z*_0_ is a scaling constant that scales free water diffusion to 1 so that the QA value will mean the same thing across different participants^13^.

Figure 2 shows that QA can be defined for each peak in the SDF. Because tractography follows individual peaks across a string of voxels, researchers typically have focused on the first peak (*QA*_0_), and have additionally normalized the *QA*_0_ metric so that it can be compared across different participants. This normalized QA metric, nQA, is calculated according to the generalized q-sampling imaging method from Yeh et al.^13^.

**Figure 2.**
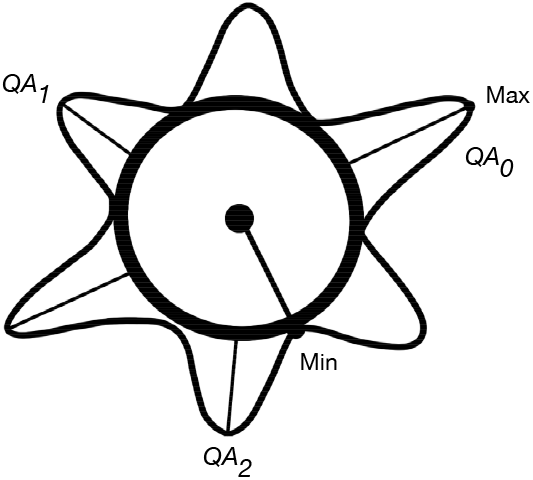
The spin distribution function (SDF) is illustrated. Each peak in QA determines a diffusion direction, with *QA*_0_ representing the primary direction, *QA*_1_ representing the secondary direction, and *QA*_2_ representing the tertiary direction.

##### Generalized Fractional Anisotropy (GFA)

Generalized fractional anisotropy (GFA) represents an alternative indirect metric of white matter integrity that can be computed from a HARDI diffusion acquisition. It can be thought of as a higher-order generalization of FA^28^. The GFA metric thus begins with the ODF, and proceeds to rescale it by subtracting off the baseline term. Rescaling the ODF introduces a confound such that noise in the data, which rescales non-linearly, can appear to be anisotropic when in fact that is not the case. The GFA corrects for this by rescaling the min-max normalized ODF with an anisotropy measure. From Tuch^16^, the GFA metric follows the same logic as the FA calculation. Thus:

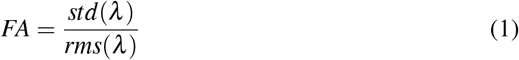

where *λ* are the eigenvalues of the diffusion tensor.

GFA is then given by:

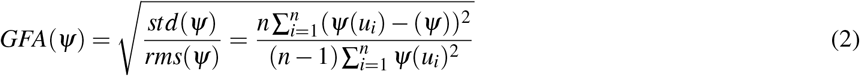

where 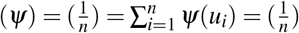 is the mean of the ODF. The output of this transformation is a scalar measure, GFA, which functions in a similar manner as FA to describe the anisotropic direction of water diffusion in the voxel. Like the traditional FA metric from DTI, the values range from 0 to 1.

##### Neurite orientation dispersion and density imaging (NODDI) Metrics

NODDI works by combining the three-component tissue model, which distinguishes between intracellular, extracellular, and cerebrospinal fluid, with the 2-shell HARDI protocol:

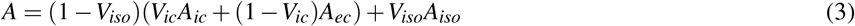

This calculates the independent, normalized signal *A*, comprised of the intra-cellular normalized signal and volume fraction (*V_ic_* and *A_ic_*), the normalized signal of the extracellular compartment (*A_ec_*), and the normalized signal and and volume fraction of the cerebrospinal fluid (*V_iso_* and *A_iso_*)^2^. The three tissue components are calculated from a simplified form of the orientation-dispersed cylinder model^29^ and the Watson distribution. Using the Watson distribution provides a unique advantage due to its capability of accurately estimating both low and high levels of dispersion across the brain, while truncated spherical harmonic series tend to provide inexact measurements of lower levels of orientation dispersion^2, 30^.

NODDI supports the modelling of both gray and white matter, with white matter showing low to moderate axonal dispersion and gray matter showing high axonal dispersion. Since its implementation, NODDI has been used in numerous adult studies to examine disease progression^31, 32^, and it appears to be useful for differentiating between myelinated and non-myelinated axons^33^ and measuring cortical gray matter maturation over development^34^.

The three NODDI metrics we examined were intracellular volume fraction (ICVF), isotropic volume fraction (ISOVF), and orientation dispersion (OD). ICVF measures neurite density, with higher values indicating greater packing of neuronal tissue. ISOVF measures the extracellular, free water compartment of the model. OD measures dispersion of modelled sticks, with greater dispersion seen in gray matter and lower dispersion in white matter regions such as the corpus callosum.

##### Restricted Diffusion Imaging (RDI)

Restricted diffusion imaging (RDI) is a novel metric that aims to measure changes in cellular density. Thus far it has been shown to be sensitive to the inflammatory response (macrophage infiltration) in rats^1^. It is for this reason that we expect it might be sensitive to differences in axonal density.

RDI works through the use of q-ball imaging that estimates the density of diffusing spins with respect to their diffusion displacement. RDI separates non-restricted diffusion from restricted diffusion through the use of different diffusion sensitization strengths, which allows RDI to be more sensitive to structural changes. The calculation for the metric is a linear combination of diffusion-weighted imaging signals acquired by the long diffusion time^1^;

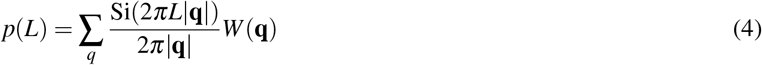

In the formula, rho (*L*) represents the density of diffusing spins that are restricted with the displacement distance *L*. Si is a sine integral, 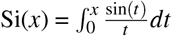. In the case where **q** = 0, the term Si(2*πL*|**q**|)/2*π*|**q**| would be replaced by *L*. Overall restricted diffusion, *ρ*(*L*), estimates the density of diffusing spins with diffusion displacements less than *L*^1^.

### DWI Reconstruction

DTI^26^ and GQI^13^ reconstruction were both done using the January 2018 version of DSIStudio (http://dsi-studio.labsolver.org) and took approximately one minute for DTI and two minutes for GQI. NODDI reconstruction took considerably longer, at about four to six hours per brain. The NODDI reconstruction done here used the AMICO^35^ python implementation, along with the DIPY library^36^, SPArse Modeling Software (SPAMS)^37^ (http://spams-devel.gforge.inria.fr/), and the Camino toolkit as dependencies^38^.

#### Parcellating the Corpus Callosum: Drawing the Regions of Interest (ROIs)

The corpus callosum was manually segmented into 10 ROIs in DSI Studio, following the segmentation scheme described by Aboitiz et al^18^. These divisions are shown in Figure 3A, reconstructed from the original Aboitiz et al (1992) segmentation. To conduct the segmentation, we first drew the midline slice from the coronal view, and then we drew the ROIs from the sagittal view. We then measured the corpus callosum divided it into three equal length sections going from the anterior to posterior direction: the genu, the midbody, and the isthmus/splenium. We then further divided the genu and the midbody sections into three equal parts. Thus, the genu was divided into G1, G2, and G3, while the midbody was divided into B1, B2, and B3.

**Figure 3.**
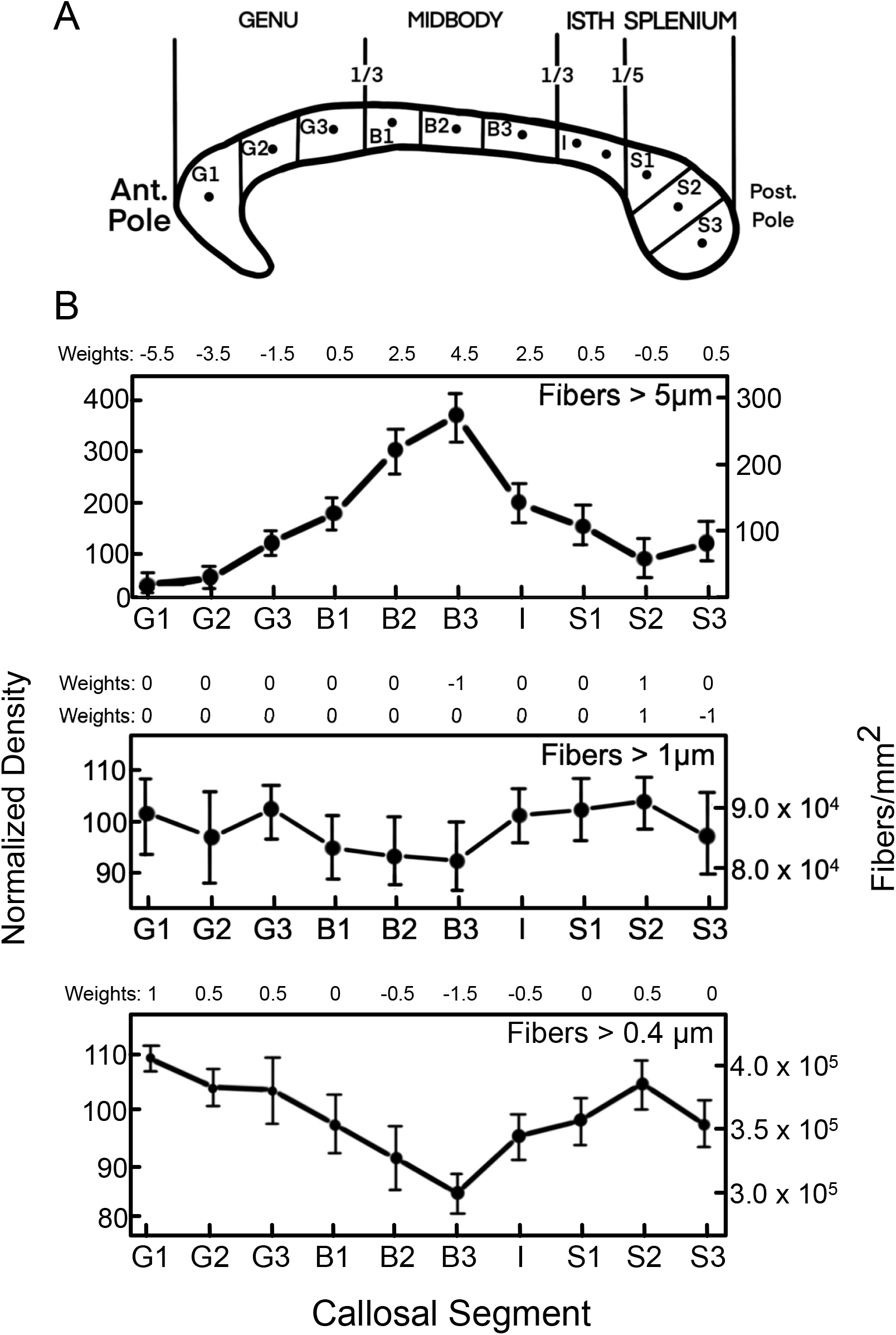
Figure 3A: Corpus callosum segmentation scheme from the Aboitiz et al (1992) paper which we replicated for the current studies. Figure 3B: Density results from Aboitiz et al (1992) for three fiber sizes: >5.0 *μ*m, >1 *μ*m, and >0.4 *μ*m, along with the associated contrast weights we applied.

Finally, we defined the last third of the corpus callosum as the isthmus and the splenium. The splenium makes up the posterior fifth of the entire corpus callosum and is subdivided into three equal sections, labelled S1, S2, and S3. The isthmus, labelled I, is comprised of the remaining area between the midbody and the splenium sections. All sections of the corpus callosum are arbitrarily divided based on straight length by counting voxels between the parcels. Clarke^39^ showed that whether these sections are portioned based on curvature or straight length makes no difference. For this adult sample, the 10 ROIs were saved as an atlas which was then applied to all participants.

#### Data Analysis

In the original Aboitiz et al., (1992) paper, the authors used this segmentation to report on densities, based on histological counts, of the following fiber sizes: >0.4 *μ*m, >1 *μ*m, >3 *μ*m, and >5.0 *μ*m. The pattern of fiber densities was nearly identical for the >3 *μ*m B and >5.0 *μ*m, and thus for simplicity we ignored that pattern for this paper (i.e., the >5.0 *μ*m results would apply in the same way to the >3.0 *μ*m pattern). In order to determine which of the density patterns produced by our two samples best replicated the Aboitiz histological model, we conducted planned contrast analyses^40^. Planned contrasts allow a test of whether the observed pattern matches an expected pattern, which in this case is defined by the histologic results.

The patterns we modeled are given in Figure 3. Figure 3A shows the segmentation scheme used in the Aboitiz et al (1992) paper, which we replicated here. This scheme was also used by Genc et al^22^. Figure 3B shows density results from Aboitiz et al (1992) for three fiber sizes: >5.0 *μ*m, >1 *μ*m, and >0.4 *μ*m.

The first step in a planned contrast analysis is to establish the contrast weights. These weights are designed to model the pattern of the Aboitiz data for the four comparisons on which they reported (again, note that results for >3 *μ*m are omitted). Figure 3B shows the contrast weights for each of four comparisons that were reported as significant in the Aboitiz paper. For >5.0 *μ*m the contrast weights were: (−5.5, −3.5, −1.5, 0.5, 2.5, 4.5, 2.5, 0.5, −0.5, 0.5). For >1 *μ*m, Aboitiz only found significant differences between B3 and S2 segments and S2 and S3 segments. Therefore we created two contrasts to test those specific differences: >1 *μ*m A: (0, 0, 0, 0, 0, −1, 0, 0, 1, 0); >1 *μ*m B: (0, 0, 0, 0, 0, 0, 0, 0, 1, −1). Finally, for >0.4 *μ*m the weights were: (1, 0.5, 0.5, 0, −0.5, −1.5, −0.5, 0, 0.5, 0). As is standard, all contrast weights sum to zero. Once contrast weights were established, the second step was to test them in the context of the linear model, which was done using R v3.6^41^.

#### Outlier Detection and Correction

We did not remove outliers but corrected for their influence using a conservative 97.5% Winsorization procedure. Similar to clipping in signal processing, this statistical transformation limits extreme values in order to reduce the influence of outliers.

**Table 1.** This summary table provides a breakdown of the metrics that were analyzed for each study. Given the long computation time, a random sample of 100 out of 840 were chosen for NODDI analyses in Study 1. Furthermore, NODDI is only available for multi-shell acquisitions, so it could not be applied to the single-shell data in Study 2.

| Metrics | *n* | Measure |
| --- | --- | --- |
| **Study 1: HCP Adults** |  |  |
| multi-shell data, *b* values = 500, 1000, 2000, 90 directions per shell |  |  |
| <u>DTI Metrics</u> |  |  |
|  | 840 | Radial Diffusivity (RD) |
|  | 840 | Axial Diffusivity (AD) |
|  | 840 | Mean Diffusivity (MD) |
|  | 840 | Fractional Anisotropy (FA) |
| <u>GQI Metrics</u> |  |  |
|  | 840 | Generalized Fractional Anisotropy (GFA) |
|  | 840 | Normalized Quantitative Anisotropy (NQA) |
| <u>NODDI Metrics</u> |  |  |
|  | 100 | Intracellular Volume Fraction (ICVF) |
|  | 100 | Isotropic Volume Fraction (ISOVF) |
|  | 100 | Orientation Dispersion (OD) |
| <u>RDI Metric</u> |  |  |
|  | 840 | Restricted Diffusion Imaging (RDI) |
| **Study 2: CMIND Children and Adolescents** |  |  |
| single shell data, *b* value = 3000, 61 directions |  |  |
| <u>DTI Metrics</u> |  |  |
|  | 129 | Radial Diffusivity (RD) |
|  | 129 | Axial Diffusivity (AD) |
|  | 129 | Mean Diffusivity (MD) |
|  | 129 | Fractional Anisotropy (FA) |
| <u>GQI Metrics</u> |  |  |
|  | 129 | Generalized Fractional Anisotropy (GFA) |
|  | 129 | Normalized Quantitative Anisotropy (NQA) |
| <u>RDI Metric</u> |  |  |
|  | 129 | Restricted Diffusion Imaging (RDI) |

#### Results

Given the large sample size, even small effects are likely to be statistically significant. Therefore, our reporting of results focuses on effect sizes, which are independent of sample size. However, in organizing the discussion of results it is useful to establish a cutoff on which to base the reporting of the most important results. We established an arbitrary cutoff of > |2.5| standard deviations to organize the discussion of the results. We still report all results in the Tables, regardless of the cutoff values. We also conducted a permutation-based significance tests (5000 iterations) to establish what is the minimum statistically significant difference in effect sizes at *α* = .05. This critical difference was 0.35.

As Table 2 and Figure 4 show, based on this cutoff, several measures were sensitive to the longitudinal distribution of fiber sizes in the corpus callosum. From the DTI metrics, AD most closely replicated histological density patterns for both the smallest fiber sizes (*t* = −60.50, *p* <0.001) and the largest fiber sizes (*t* = 70.50, *p* <0.001). From the GQI metrics, GFA was the most sensitive to fibers greater than 1 *μ*m for both contrasts: B3 and S2 (*t* = 52.42, *p* <0.001) and S2 and S3 (*t* =22.85, *p* <0.001).

**Table 2.** Contrast analyses for corpus callosum density patterns in the adult sample. Effect size reflects absolute value. *p*<0.05*, *p* <0.01**, *p* <0.001***.

| Metric | *n* | *T*-value | Cohen's *d* | *p* -value |
| --- | --- | --- | --- | --- |
| **>0.4 $\mu\text{m}$ Fiber Size** |  |  |  |  |
| Fractional Anisotropy (FA) | 840 | 9.70 | 0.47 | <0.001*** |
| Radial Diffusivity (RD) | 840 | -28.48 | 1.39 | <0.001*** |
| Axial Diffusivity (AD) | 840 | -60.60 | 2.96 | <0.001*** |
| Mean Diffusivity (MD) | 840 | -41.90 | 2.05 | <0.001*** |
| Generalized Fractional Anisotropy (GFA) | 840 | 7.18 | 0.35 | <0.001*** |
| Normalized Quantitative Anisotropy (NQA) | 840 | -19.92 | 0.97 | <0.001*** |
| Intracellular Volume Fraction (ICVF) | 100 | 25.54 | 3.61 | <0.001*** |
| Isotropic Volume Fraction (ISOVF) | 100 | -4.56 | 0.65 | <0.001*** |
| Orientation Dispersion (OD) | 100 | -0.06 | 0.01 | 0.95 |
| Restricted Diffusion Imaging (RDI) | 840 | -53.29 | 2.60 | <0.001*** |
| **>1 $\mu\text{m}$ (A) Fiber Size Comparing B3 and S2** |  |  |  |  |
| Fractional Anisotropy (FA) | 840 | 48.51 | 2.37 | <0.001*** |
| Radial Diffusivity (RD) | 840 | -39.68 | 1.94 | <0.001*** |
| Axial Diffusivity (AD) | 840 | -6.27 | 0.31 | <0.001*** |
| Mean Diffusivity (MD) | 840 | -34.47 | 1.68 | <0.001*** |
| Generalized Fractional Anisotropy (GFA) | 840 | 52.42 | 2.56 | <0.001*** |
| Normalized Quantitative Anisotropy (NQA) | 840 | 32.28 | 1.58 | <0.001*** |
| Intracellular Volume Fraction (ICVF) | 100 | 17.35 | 2.45 | <0.001*** |
| Isotropic Volume Fraction (ISOVF) | 100 | -7.71 | 1.09 | <0.001*** |
| Orientation Dispersion (OD) | 100 | -7.28 | 1.03 | <0.001*** |
| Restricted Diffusion Imaging (RDI) | 840 | -22.92 | 1.12 | <0.001*** |
| **>1 $\mu\text{m}$ (B) Fiber Size Comparing S2 and S3** |  |  |  |  |
| Fractional Anisotropy (FA) | 840 | 17.53 | 0.86 | <0.001*** |
| Radial Diffusivity (RD) | 840 | -13.99 | 0.68 | <0.001*** |
| Axial Diffusivity (AD) | 840 | 4.29 | 0.21 | <0.001*** |
| Mean Diffusivity (MD) | 840 | -10.88 | 0.53 | <0.001*** |
| Generalized Fractional Anisotropy (GFA) | 840 | 22.85 | 1.12 | <0.001*** |
| Normalized Quantitative Anisotropy (NQA) | 840 | 5.30 | 0.26 | <0.001*** |
| Intracellular Volume Fraction (ICVF) | 100 | -6.28 | 0.89 | <0.001*** |
| Isotropic Volume Fraction (ISOVF) | 100 | -9.73 | 1.38 | <0.001*** |
| Orientation Dispersion (OD) | 100 | -12.09 | 1.71 | <0.001*** |
| Restricted Diffusion Imaging (RDI) | 840 | -6.02 | 0.29 | <0.001*** |
| **>5 $\mu\text{m}$ Fiber Size** |  |  |  |  |
| Fractional Anisotropy (FA) | 840 | 4.73 | 0.23 | <0.001*** |
| Radial Diffusivity (RD) | 840 | 20.65 | 1.01 | <0.001*** |
| Axial Diffusivity (AD) | 840 | 70.50 | 3.44 | <0.001*** |
| Mean Diffusivity (MD) | 840 | 38.56 | 1.88 | <0.001*** |
| Generalized Fractional Anisotropy (GFA) | 840 | 7.83 | 0.38 | <0.001*** |
| Normalized Quantitative Anisotropy (NQA) | 840 | 35.17 | 1.72 | <0.001*** |
| Intracellular Volume Fraction (ICVF) | 100 | -24.66 | 3.49 | <0.001*** |
| Isotropic Volume Fraction (ISOVF) | 100 | 3.14 | 0.44 | <0.001*** |
| Orientation Dispersion (OD) | 100 | -1.21 | 0.17 | 0.23 |
| Restricted Diffusion Imaging (RDI) | 840 | 58.24 | 2.84 | <0.001*** |

**Figure 4.**
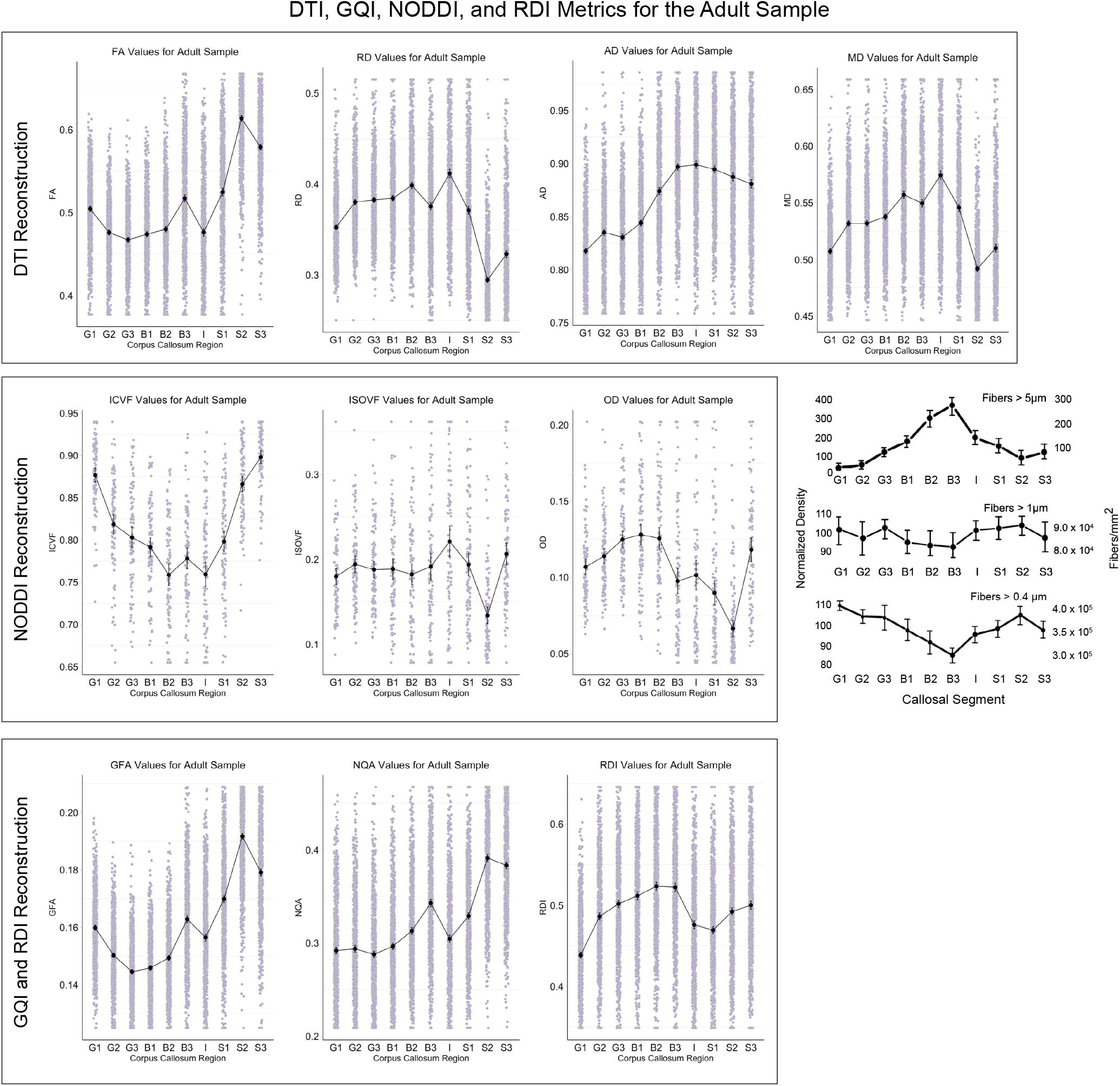
Anterior to posterior corpus callosum density patterns for the adult sample are plotted for each metric and grouped by reconstruction method. The density patterns acquired in the *in vivo* DWI acquistions are being compared to the Aboitiz (1992) histological patterns, shown on the right.

As expected, the most accurate NODDI metric was ICVF, which accurately detected density patterns at the smallest fiber size (*t* = 25.54, *p* <0.001) and at the largest fiber size (*t* = −24.66, *p* <0.001). The NODDI density patterns seen in this adult sample replicate the patterns reported by Genc et al^22^ in a developing sample. Across both studies, ICVF was the lowest at the isthmus and the highest at the posterior part of the splenium. Similarly, the novel metric reported here, RDI, accurately replicated the density pattern for the > 5 *μ*m fiber sizes (*t* = 58.24, *p* <0.001), and for the smallest, > 0.4 *μ*m fiber sizes (*t* = −53.29, *p* <0.001). These findings indicate that both RDI and NODDI are capable of accurately measuring axonal density for large and small fiber sizes. At > 0.4 and 5 *μ*m fiber sizes NODDI was statistically better than RDI, which was itself better than all other metrics except AD.

## Study 2

In Study 2 we aimed replicate the results from Study 1 in a developing sample. The same DTI and GQI diffusion metrics and histological model were used, but with a manually-drawn corpus callosum segmentation for each participant as previously detailed. NODDI metrics were not able to be calculated on this sample due to NODDI’s multi-shell requirement, not met by this sample’s single-shell MRI data. Note that Genc and colleagues^22^ did examine NODDI metrics using a second acquisition in the CMIND data set, which had different acquisition parameters and a different shell. We focused on the single shell acquisition, as we were interested in determining whether RDI can provide accurate results with one shell acquisition. However, because we used the same segmentation approach as they did, we can compare our findings to theirs in a partial replication.

### Participants

The developing sample contained 129 participants between the ages of 7-months-old and 19-years-old (*M* = 8.8 years) from the Cincinnati MR Imaging of NeuroDevelopment (C-MIND) database, provided by the Pediatric Functional Neuroimaging Research Network (https://research.cchmc.org/c-mind/) and supported by a contract from the Eunice Kennedy Shriver National Institute of Child Health and Human Development (HHSN275200900018C). The data are available from CMIND by request, which facilitates validation of the results we report here. Participants in the database are full-term gestation, healthy, righthanded, native English speakers, without contraindication to MRI. By design, the C-MIND cohort is demographically diverse (38% nonwhite, 55% female, median household income $42,500), intended to reflect the US population. The summary statistics for the DTI and GQI metrics were calculated for the full sample of 129 participants.

### Parcellating the Corpus Callosum

Due to previous studies indicating the problems with applying the MNI normalization to a developing sample, we manually drew the 10 ROIs for the Study 2 sample, following the same guidelines as were used in Study 1. The same data analysis procedure was followed as in Study 1. As in Study 1, we also conducted a permutation-based significance tests (5000 iterations) to establish what is the minimum statistically significant difference in effect sizes at α = .05. For the child sample, this critical difference was also 0.35.

### Results

There were notable differences when the diffusion metrics were applied to the C-MIND developing sample. First, AD was no longer the stand-out DTI metric. Instead, FA was the most effective DTI metric at replicating density patterns for 3 out of 4 models. It was positively related to smaller fiber sizes (*t* = 28.66, *p* <0.001 for > 0.4 *μ*m, *t* = 23.88, *p* <0.001 for > 1 *μ*m model A) and was inversely related to the largest > 5 *μ*m density model (*t* = −27.46, *p* <0.001).

For the GQI metrics, GFA continued to be a reliable measure of fiber density for all four models. RDI was also consistent for both the smallest and highest fiber size models. In fact, RDI most closely matched the histological model for fiber sizes greater than 5 *μ*m (*t* = 30.36, *p* <0.001), and was statistically better than all other metrics.

**Figure 5.**
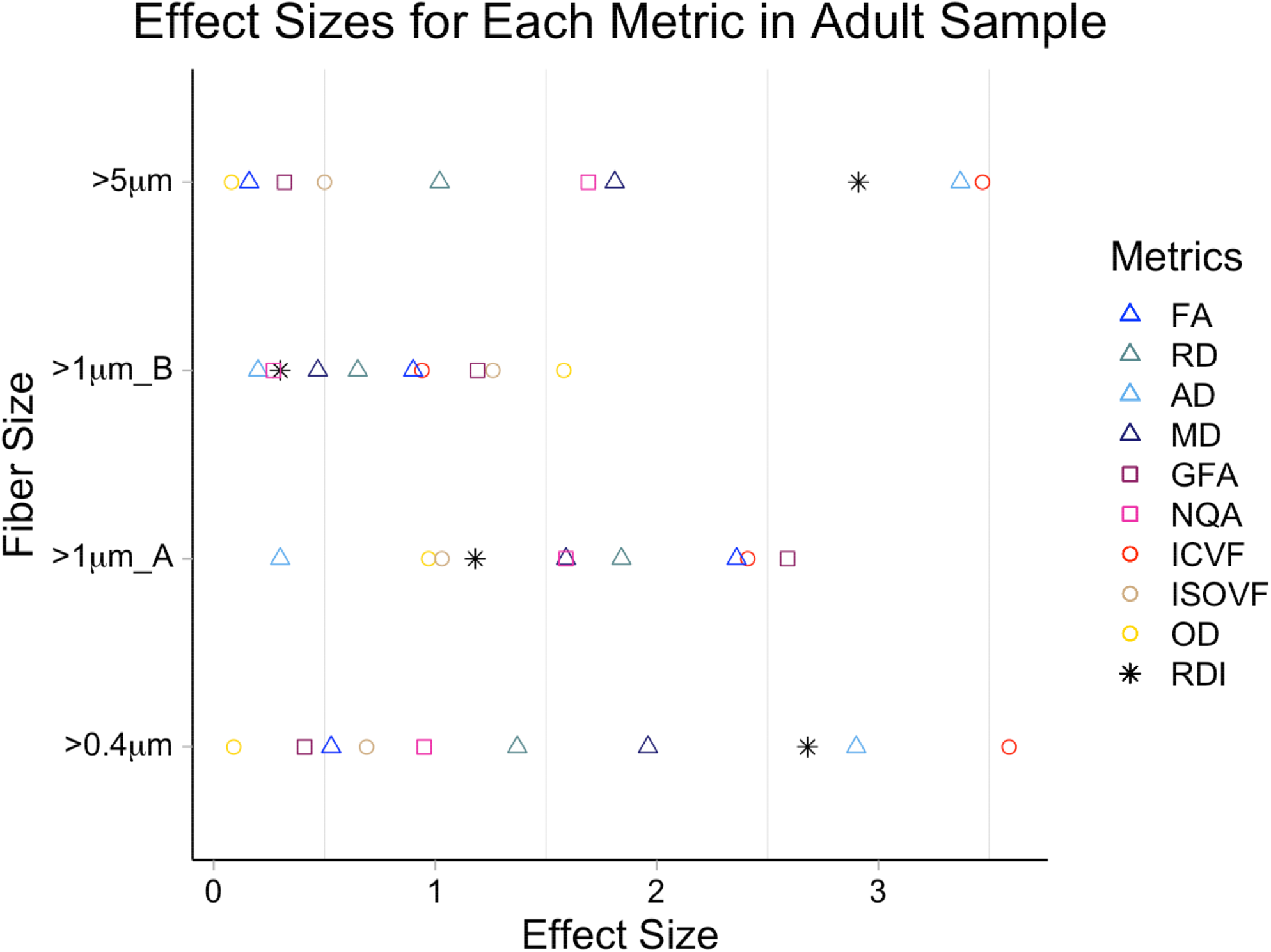
This figure shows the effect size of each diffusion metric predicting the Aboitiz model in the adult sample, based on fiber size. Absolute value for effect size is displayed. Fiber sizes above 1*μ*M compared the differences in B3 and S2 for model A, and differences between fiber sizes greater than 1*μ*M in S2 and S3 for model B.

**Figure 6.**
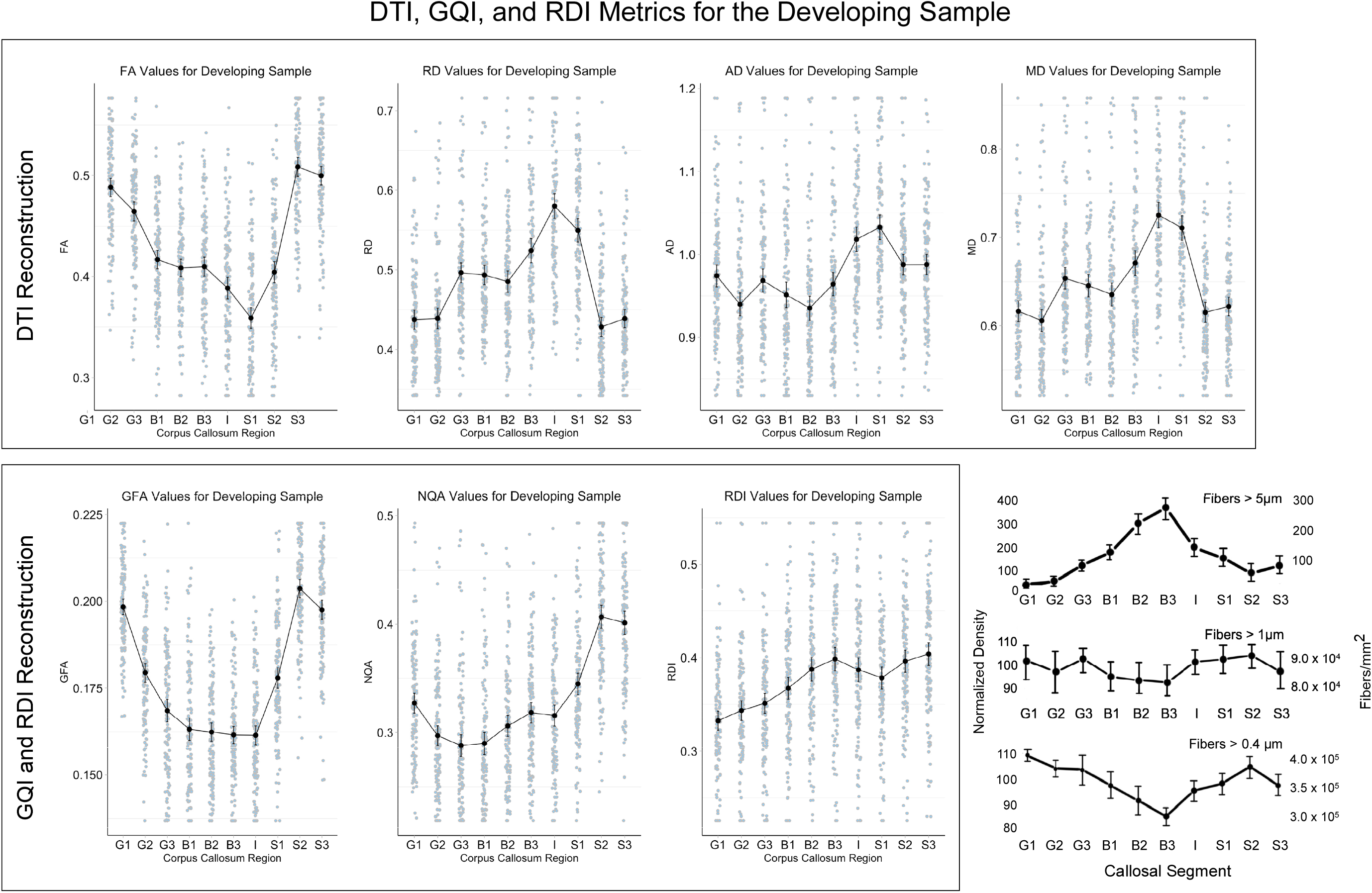
Corpus callosum density patterns for the developing sample are plotted for each of the 7 metrics and grouped by reconstruction method. The density patterns acquired in the *in vivo* DWI acquistions are being compared to the Aboitiz (1992) histological patterns, shown on the bottom right.

**Figure 7.**
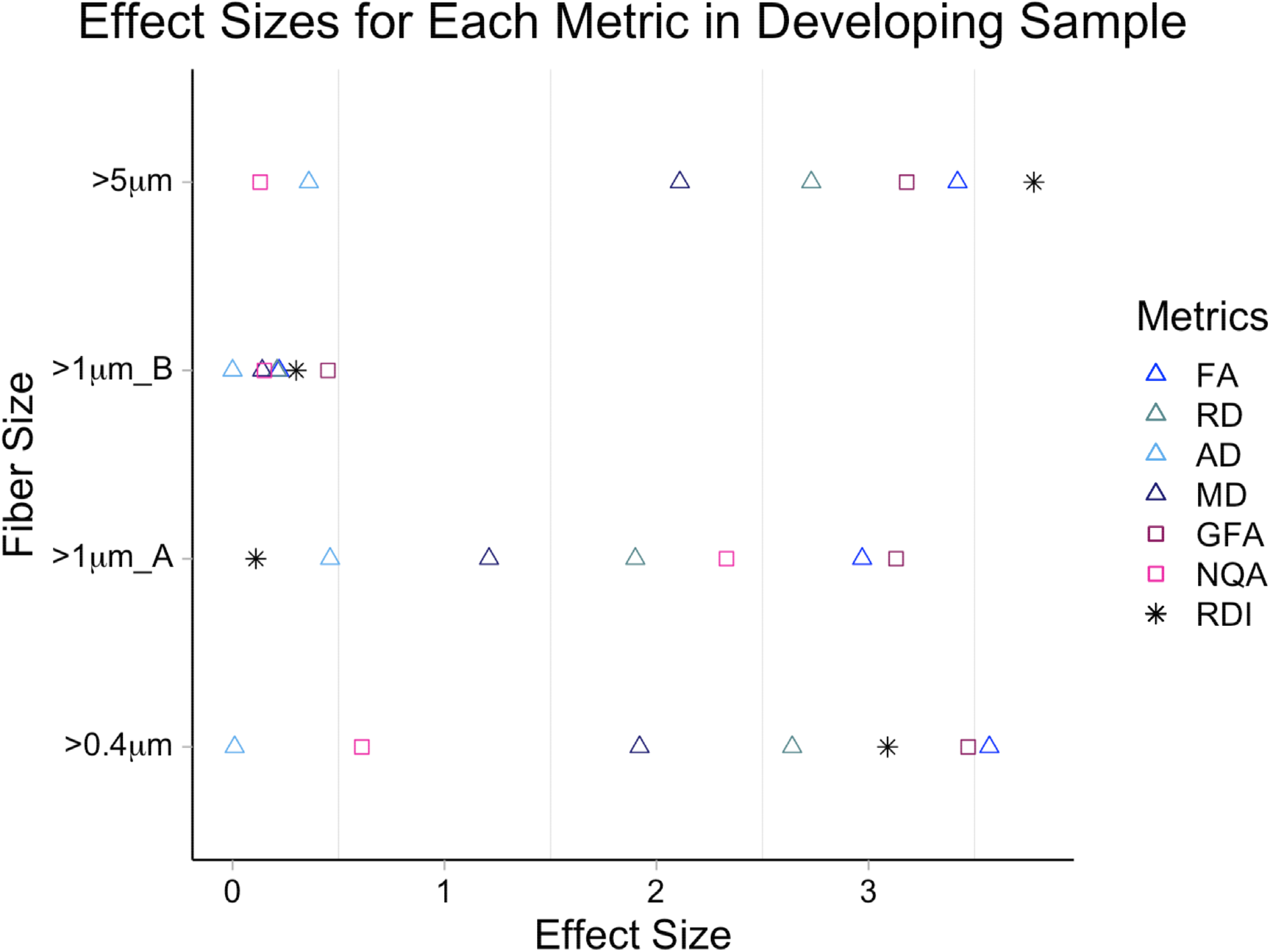
This figure shows the effect size of each diffusion metric predicting the Aboitiz model in the developing sample, based on fiber size. Absolute value of effect size is displayed. Fiber sizes above 1*μ*M compared the differences in B3 and S2 for model A, and differences between fiber sizes greater than 1*μ*M in S2 and S3 for model B.

**Table 3.** Contrast analyses for corpus callosum density patterns in the developing sample. Effect size reflects absolute value. *p* <0.05*, *p* <0.01**, *p* <0.001***.

| Metric | *n* | *T*-value | Cohen's *d* | *p* -value |
| --- | --- | --- | --- | --- |
| **>0.4 $\mu\text{m}$ Fiber Size** |  |  |  |  |
| Fractional Anisotropy (FA) | 129 | 28.66 | 3.57 | <0.001*** |
| Radial Diffusivity (RD) | 129 | -21.23 | 2.64 | <0.001*** |
| Axial Diffusivity (AD) | 129 | -0.10 | 0.01 | 0.92 |
| Mean Diffusivity (MD) | 129 | -15.40 | 1.92 | <0.001*** |
| Generalized Fractional Anisotropy (GFA) | 129 | 27.86 | 3.47 | <0.001*** |
| Normalized Quantitative Anisotropy (NQA) | 129 | 4.88 | 0.61 | <0.001*** |
| Restricted Diffusion Imaging (RDI) | 129 | -24.79 | 3.09 | <0.001*** |
| **>1 $\mu\text{m}$ (A) Fiber Size Comparing B3 and S2** |  |  |  |  |
| Fractional Anisotropy (FA) | 129 | 23.88 | 2.97 | <0.001*** |
| Radial Diffusivity (RD) | 129 | -15.29 | 1.90 | <0.001*** |
| Axial Diffusivity (AD) | 129 | 3.67 | 0.46 | <0.001*** |
| Mean Diffusivity (MD) | 129 | -9.71 | 1.21 | <0.001*** |
| Generalized Fractional Anisotropy (GFA) | 129 | 25.16 | 3.13 | <0.001*** |
| Normalized Quantitative Anisotropy (NQA) | 129 | 18.70 | 2.33 | <0.001*** |
| Restricted Diffusion Imaging (RDI) | 129 | -0.85 | 0.11 | 0.40 |
| **>1 $\mu\text{m}$ (B) Fiber Size Comparing S2 and S3** |  |  |  |  |
| Fractional Anisotropy (FA) | 129 | 1.15 | 0.22 | 0.08 |
| Radial Diffusivity (RD) | 129 | -1.67 | 0.21 | 0.10 |
| Axial Diffusivity (AD) | 129 | -0.02 | 0.003 | 0.98 |
| Mean Diffusivity (MD) | 129 | -1.16 | 0.14 | 0.24 |
| Generalized Fractional Anisotropy (GFA) | 129 | 3.64 | 0.45 | <0.001*** |
| Normalized Quantitative Anisotropy (NQA) | 129 | 1.17 | 0.17 | 0.24 |
| Restricted Diffusion Imaging (RDI) | 129 | -2.44 | 0.30 | <0.05* |
| **>5 $\mu\text{m}$ Fiber Size** |  |  |  |  |
| Fractional Anisotropy (FA) | 129 | -27.46 | 3.42 | <0.001*** |
| Radial Diffusivity (RD) | 129 | 21.94 | 2.73 | <0.001*** |
| Axial Diffusivity (AD) | 129 | 2.87 | 0.36 | <0.01** |
| Mean Diffusivity (MD) | 129 | 16.93 | 2.11 | <0.001*** |
| Generalized Fractional Anisotropy (GFA) | 129 | -25.55 | 3.18 | <0.001*** |
| Normalized Quantitative Anisotropy (NQA) | 129 | 1.04 | 0.13 | 0.30 |
| Restricted Diffusion Imaging (RDI) | 129 | 30.36 | 3.78 | <0.001*** |

## Discussion

The quantification of microstructural properties of neural tissue can be informed by advances in the modeling of diffusion-weighted imaging data. This is important for the diagnosis and treatment of neurologic disease, especially as it relates to properties of white matter such as axon density. In this study, we examined two separate samples of typical adults and children. In the first sample of 840 adults from the HCP dataset, we segmented the corpus callosum into 10 parts, and measured the diffusion properties of each segment. These diffusion properties were modeled using a variety of reconstruction methods based on DTI, NODDI, and GQI models. For large fiber sizes (> 5*μ*m), we found that the best indicators of histologically established axonal density patterns were the DTI AD, NODDI ICVF, and RDI. For mid-range fiber sizes (> 1*μ*m), there were no methods that were clear standouts. For the smallest fiber sizes (> 0.4*μ*m), the same metrics that were sensitive to large fiber sizes (DTI AD, NODDI ICVF, and RDI) showed the clearest differentiation.

For the developing sample of 129 children and adolescents, we generally found the same results as in the adult sample, but there were a couple of exceptions. Unlike the adult sample, DTI AD was not a clear differentiator. Additional measures (i.e., DTI FA and GQI GFA) were also sensitive measures, but this was not evident in the adult sample. We did, however, replicate the finding for the RDI metric. As in the adult sample, the RDI metric clearly matched the known histologic pattern of the corpus callosum, especially for small and large fiber sizes, and indeed seemed to be sensitive to subtle differences across the longitudinal length of the structure. These results suggest it is possible to quantify, indirectly, differences in axonal density *in vivo* using DWI in both children and adults. Furthermore, there is clear support that RDI is a useful metric of fiber density in both adults and children, and in single- and multi-shell acquisition paradigms. This information has the potential to inform researchers and clinicians about microstructural differences in a variety of domains, for example in the examination of disease progression, of age-related changes across typical and atypical development, and of general nervous system structure/function relationships.

### Comparison of Restricted Diffusion Imaging (RDI) to Other Metrics in Published Literature

Our study can be conceived of as a complement to similar investigations conducted by Genc and colleagues^22^ and by Raffelt and colleagues^42, 43^. These authors showed that other metrics sensitive to fiber density can reliably replicate the longitudinal pattern of axonal density in the corpus callosum. For example, Raffelt and colleagues^42, 43^ established the apparent fiber density (AFD) measure, which they showed can be biologically interpreted as a measure of the intra-axonal volume fraction of axons along the corresponding orientation. However, the metric is heavily affected by the space occupied by axons, and not so much by axonal diameter. Thus, for example, fiber bundles consisting of large axons could have lower density than thinner axons, yet their AFD would be comparable. Despite this, Genc and colleagues^22^ showed that AFD is sensitive to axonal density in larger fiber bundles (their Figure 4), and indeed that metric best matched the histological pattern reported by Aboitiz. But Genc et al., did not assess whether it was similarly sensitive for smaller fiber bundles.

In our complementary investigation, we showed that, for both large and small fiber sizes, in both adult and child samples, RDI was an excellent differentiator of the pattern of axonal density across the longitudinal axis of the corpus callosum (see Figure 8 for comparison with large diameter fibers). Unlike most other diffusion metrics, RDI does does not depend on an underlying diffusion model or numerical optimization to estimate model parameters, making it easily applicable to a wider range of databases and clinical scanner protocols. RDI’s sensitivity to cell density has been useful in differentiating tissue in patients with tumors^44, 45^, as well as measuring therapeutic benefits of deep brain stimulation^46^. In a phantom study by Yeh et al, the optimized restricted diffusion showed an almost perfect correlation of 0.998 with cell density^1^. Notably, that paper also showed that mean diffusivity is associated with cell density. However, we did not corroborate that here with respect to axonal density. That is, MD is not as good as RDI at differentiating the longitudinal pattern of fiber density in the corpus callosum. Thus, despite similar sensitivity to cellular density in a phantom study, RDI seems to be more sensitive than MD to fiber density *in vivo*. Furthermore, RDI was highly consistent across the two samples (see Figure 8). This is in contrast to other common metrics like FA, which was not consistent across the two samples. These results suggest that RDI is a reliable measure of fiber density across a wide age range, and it is robust to different diffusion acquisitions. Furthermore, a main advantage of RDI is the computation time (under a minute) which occurs as part of routine image reconstruction, thus making it appealing in clinical settings.

**Figure 8.**
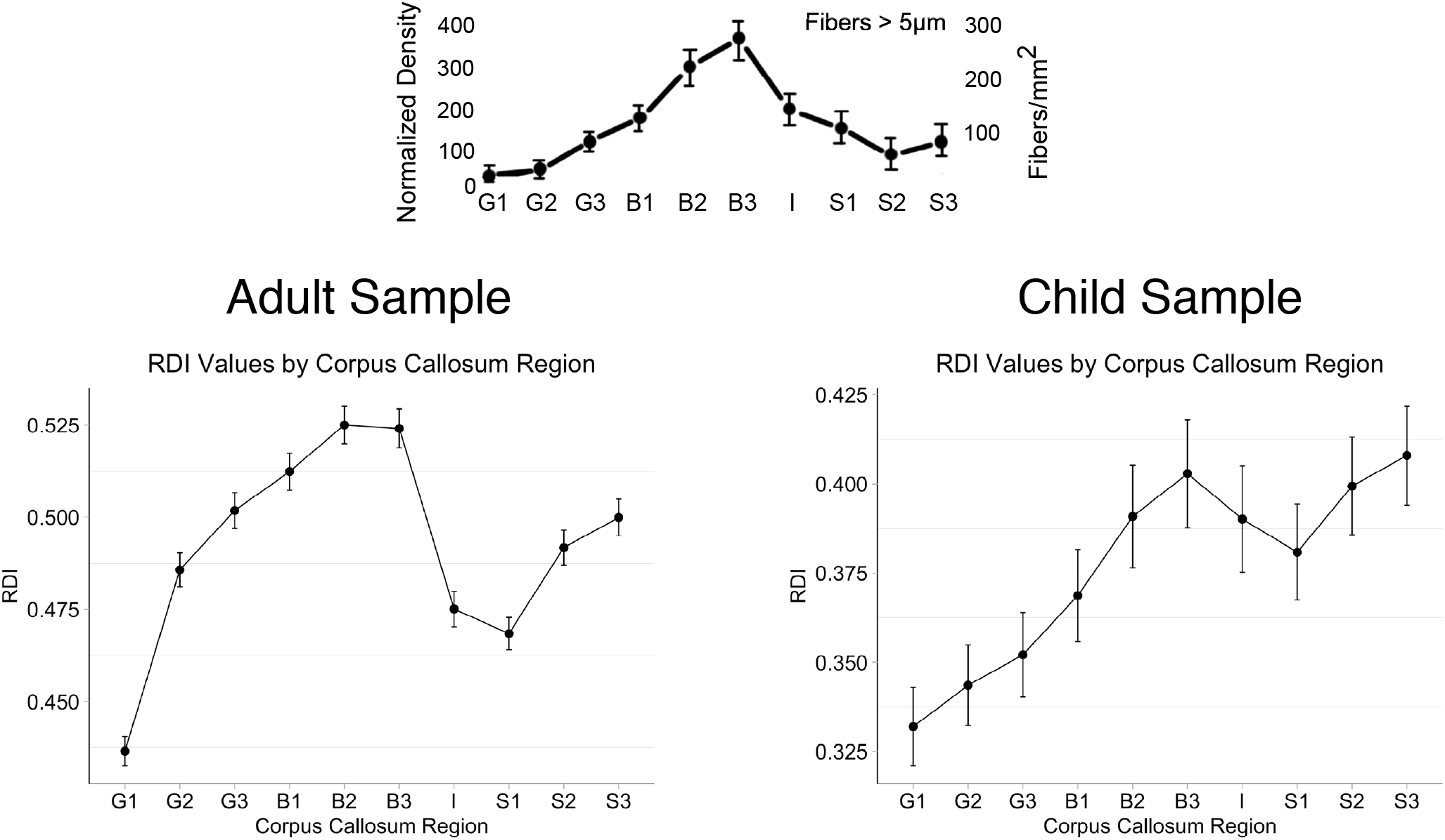
The Aboitiz (1992) histological density model for large fiber sizes is shown above. The bottom plots illustrate the density patterns for the adult and child samples as measured by RDI. RDI accurately captures the peaks and troughs of histologically-established axonal density patterns along the longitudinal axis of the corpus callosum.

Genc and colleagues^22^ also showed that NODDI metrics were reasonably sensitive to the distribution of axon density in the corpus callosum. In our study, for large fiber sizes, NODDI ICVF was the best NODDI differentiator of corpus callosum fiber density. In other research, this measure has also been shown to be effective at capturing axonal density across the corpus callosum^2, 22^. For example, previous studies have found that NODDI is more sensitive to demyelination than DTI metrics in both human^47^ and mouse models^48^. However, despite the sensitivity, two issues limit the utility of NODDI for clinical samples, especially for pediatric samples. One is the long reconstruction times (several hours per participant in our study), and the second is the requirement of multi-shell data acquisitions. Some software advances have improved reconstruction times for NODDI models (to around 1 hour per participant using Microstructure Diffusion Toolbox (MDT)^49^), but RDI reconstruction times are still much shorter (around 1 minute per participant), and our data suggest that, for fiber density, RDI can provide accurate results even for single-shell acquisitions. This does not entirely discount the effectiveness of NODDI ICVF as a metric for assessing fiber density, but it does suggest that RDI is another useful addition to the diffusion toolkit, and that it has important clinical applicability.

RDI also outperformed DTI metrics. For our adult sample, our density patterns were similar to previous investigations, with an increase in FA seen in the posterior segments of the corpus callosum^20, 21^ and a peak of MD in the isthmus^21^, indicating that DTI metrics are sensitive to structural differences in the corpus callosum. But DTI metrics were inconsistent between the adult and child samples. Furthermore, despite being widely utilized as measures of microstructural change, DTI metrics lack specificity when it comes to measuring specific characteristics of microstructure^14^. The inconsistencies we saw in our DTI metrics might have been caused by differences in axonal density, but they might also be associated with other tissue properties, such as lower myelination^50, 51^, increased orientation dispersion^52^, or less mature axonal integrity within the developing sample^53, 54^. Due to the inherent nonspecific nature of DTI, it remains difficult to discern what caused the differences in quantification between samples, which highlights well-established drawbacks to DTI metrics.

### Limitations

One inherent limitation of this study is that there are many more available diffusion-weighted imaging reconstruction algorithms than we studied here. We restricted our comparison to the novel RDI method and compared it to DTI, GQI, and NODDI techniques, but there are other algorithms that should also be considered. A second limitation of our study is that we only examined one brain region. The corpus callosum is a coherently orientated structure with no crossing fibers. Our study does not address how accurate these metrics would be at measuring density when crossing fibers are present, which is a known drawback of DTI in particular, but might also affect more recently-developed metrics such as NODDI and RDI. While our study provides strong evidence for the utility of NODDI and RDI for measuring axonal density, investigations of other brain regions would further expand these results and potentially elucidate different clinical applications.

### Conclusions

The study provided novel evidence that RDI can be used to measure axonal density in children and adults, even for single-shell data. The histological model of corpus callosum anterior-posterior density pattern was replicated using this novel *in-vivo* metric across two independent samples. NODDI’s ICVF measure was also sensitive to axonal density in the adult population. The encouraging findings support the hypothesis that RDI and ICVF can be used for measurement of axonal density as a biomarker for demyelination disorders, such as multiple sclerosis, in the future.

Establishing reliable and effective measures of axonal density is potentially significant for the study of early symptomatology, progression, and possible underlying causes of disorders characterized by white matter and other neural pathologies, such as multiple sclerosis^47^, amyotrophic lateral sclerosis^32^, dysplasia^55^, epilepsy^55^, and immune cell infiltration^1^. Many previous relationships between axonal density and clinical disorders were fully dependent on histology, but through the advancements of DWI, these disorders can be studied *in vivo*, which holds potential for making earlier diagnoses and may contribute to the design of more specific, directed treatment plans.

## Acknowledgements

This research was supported by NIH Grants R01MH112588 and R01DK119814 to P.G. and A.S.D.

## References

1. Yeh, F. C., Liu, L., Hitchens, T. K. & Wu, Y. L. Mapping immune cell infiltration using restricted diffusion mri. Magn Reson. Med 77, 603–612, DOI: 10.1002/mrm.26143 (2017).

2. Zhang, H., Schneider, T., Wheeler-Kingshott, C. A. & Alexander, D. C. Noddi: practical in vivo neurite orientation dispersion and density imaging of the human brain. Neuroimage 61, 1000–16, DOI: 10.1016/j.neuroimage.2012.03.072 (2012).

3. Tallantyre, E. C. et al. Greater loss of axons in primary progressive multiple sclerosis plaques compared to secondary progressive disease. Brain 132, 1190–9, DOI: 10.1093/brain/awp106 (2009).

4. Deluca, G. C., Ebers, G. C. & Esiri, M. M. The extent of axonal loss in the long tracts in hereditary spastic paraplegia. Neuropathol Appl Neurobiol 30, 576–84, DOI: 10.1111/j.1365-2990.2004.00587.x (2004).

5. Seehusen, F. & Baumgartner, W. Axonal pathology and loss precede demyelination and accompany chronic lesions in a spontaneously occurring animal model of multiple sclerosis. Brain Pathol 20, 551–9, DOI: 10.1111/j.1750-3639.2009.00332.x (2010).

6. Ritchie, J. M. On the relation between fibre diameter and conduction velocity in myelinated nerve fibres. Proc R Soc Lond B Biol Sci 217, 29–35, DOI: 10.1098/rspb.1982.0092 (1982).

7. Song, S. K. et al. Diffusion tensor imaging detects and differentiates axon and myelin degeneration in mouse optic nerve after retinal ischemia. Neuroimage 20, 1714–22, DOI: 10.1016/j.neuroimage.2003.07.005 (2003).

8. Song, S. K. et al. Demyelination increases radial diffusivity in corpus callosum of mouse brain. Neuroimage 26, 132–40, DOI: 10.1016/j.neuroimage.2005.01.028 (2005).

9. Beaulieu, C. The basis of anisotropic water diffusion in the nervous system - a technical review. NMR Biomed 15, 435–55, DOI: 10.1002/nbm.782 (2002).

10. Chan, A. H. et al. Neural correlates of nouns and verbs in early bilinguals. Ann N Y Acad Sci 1145, 30–40, DOI: 10.1196/annals.1416.000 (2008).

11. Lebel, C. & Beaulieu, C. Lateralization of the arcuate fasciculus from childhood to adulthood and its relation to cognitive abilities in children. Hum Brain Mapp 30, 3563–73, DOI: 10.1002/hbm.20779 (2009).

12. Nir, T. M. et al. Effectiveness of regional dti measures in distinguishing alzheimer’s disease, mci, and normal aging. Neuroimage Clin 3, 180–95, DOI: 10.1016/j.nicl.2013.07.006 (2013).

13. Yeh, F. C., Wedeen, V. J. & Tseng, W. Y. Generalized q-sampling imaging. IEEE Trans Med Imaging 29, 1626–35, DOI: 10.1109/TMI.2010.2045126 (2010).

14. Daducci, A. et al. Quantitative comparison of reconstruction methods for intra-voxel fiber recovery from diffusion mri. IEEE Trans Med Imaging 33, 384–99, DOI: 10.1109/TMI.2013.2285500 (2014).

15. Cohen-Adad, J. et al. Detection of multiple pathways in the spinal cord using q-ball imaging. Neuroimage 42, 739–49, DOI: 10.1016/j.neuroimage.2008.04.243 (2008).

16. Tuch, D. S. Q-ball imaging. Magn Reson. Med 52, 1358–72, DOI: 10.1002/mrm.20279 (2004).

17. Genc, S., Malpas, C. B., Holland, S. K., Beare, R. & Silk, T. J. Neurite density index is sensitive to age related differences in the developing brain. Neuroimage 148, 373–380, DOI: 10.1016/j.neuroimage.2017.01.023 (2017).

18. Aboitiz, F., Scheibel, A. B., Fisher, R. S. & Zaidel, E. Fiber composition of the human corpus callosum. Brain Res 598, 143–53, DOI: 10.1016/0006-8993(92)90178-c (1992).

19. Suzuki, Y. et al. Estimation of the mean axon diameter and intra-axonal space volume fraction of the human corpus callosum: Diffusion q-space imaging with low q-values. Magn Reson. Med Sci 15, 83–93, DOI: 10.2463/mrms.2014-0141 (2016).

20. Caminiti, R. et al. Diameter, length, speed, and conduction delay of callosal axons in macaque monkeys and humans: comparing data from histology and magnetic resonance imaging diffusion tractography. J Neurosci 33, 14501–11, DOI: 10.1523/JNEUROSCI.0761-13.2013 (2013).

21. Bjornholm, L. et al. Structural properties of the human corpus callosum: Multimodal assessment and sex differences. Neuroimage 152, 108–118, DOI: 10.1016/j.neuroimage.2017.02.056 (2017).

22. Genc, S., Malpas, C. B., Ball, G., Silk, T. J. & Seal, M. L. Age, sex, and puberty related development of the corpus callosum: a multi-technique diffusion mri study. Brain Struct Funct 223, 2753–2765, DOI: 10.1007/s00429-018-1658-5 (2018).

23. Reyes-Haro, D., Mora-Loyola, E., Soria-Ortiz, B. & Garcia-Colunga, J. Regional density of glial cells in the rat corpus callosum. Biol Res 46, 27–32, DOI: 10.4067/S0716-97602013000100004 (2013).

24. Riise, J. & Pakkenberg, B. Stereological estimation of the total number of myelinated callosal fibers in human subjects. J Anat 218, 277–84, DOI: 10.1111/j.1469-7580.2010.01333.x (2011).

25. Wilke, M., Holland, S. K., Altaye, M. & Gaser, C. Template-o-matic: a toolbox for creating customized pediatric templates. Neuroimage 41, 903–13, DOI: 10.1016/j.neuroimage.2008.02.056 (2008).

26. Basser, P. J., Mattiello, J. & LeBihan, D. Mr diffusion tensor spectroscopy and imaging. Biophys J 66, 259–67, DOI: 10.1016/S0006-3495(94)80775-1 (1994).

27. Hasan, K. M. & Narayana, P. A. Retrospective measurement of the diffusion tensor eigenvalues from diffusion anisotropy and mean diffusivity in dti. Magn Reson. Med 56, 130–7, DOI: 10.1002/mrm.20935 (2006).

28. Descoteaux, M., Angelino, E., Fitzgibbons, S. & Deriche, R. Regularized, fast, and robust analytical q-ball imaging. Magn Reson. Med 58, 497–510, DOI: 10.1002/mrm.21277 (2007).

29. Zhang, H., Hubbard, P. L., Parker, G. J. & Alexander, D. C. Axon diameter mapping in the presence of orientation dispersion with diffusion mri. Neuroimage 56, 1301–15, DOI: 10.1016/j.neuroimage.2011.01.084 (2011).

30. Jespersen, S. N., Kroenke, C. D., Ostergaard, L., Ackerman, J. J. & Yablonskiy, D. A. Modeling dendrite density from magnetic resonance diffusion measurements. Neuroimage 34, 1473–86, DOI: 10.1016/j.neuroimage.2006.10.037 (2007).

31. Taoka, T. et al. White matter microstructural changes in tuberous sclerosis: Evaluation by neurite orientation dispersion and density imaging (noddi) and diffusion tensor images. Sci Rep 10, 436, DOI: 10.1038/s41598-019-57306-w (2020).

32. Broad, R. J. et al. Neurite orientation and dispersion density imaging (noddi) detects cortical and corticospinal tract degeneration in als. J Neurol Neurosurg Psychiatry 90, 404–411, DOI: 10.1136/jnnp-2018-318830 (2019).

33. Kunz, N. et al. Assessing white matter microstructure of the newborn with multi-shell diffusion mri and biophysical compartment models. Neuroimage 96, 288–99, DOI: 10.1016/j.neuroimage.2014.03.057 (2014).

34. Pecheva, D. et al. Recent advances in diffusion neuroimaging: applications in the developing preterm brain. F1000Res 7, DOI: 10.12688/f1000research.15073.1 (2018).

35. Daducci, A. et al. Accelerated microstructure imaging via convex optimization (amico) from diffusion mri data. Neuroimage 105, 32–44, DOI: 10.1016/j.neuroimage.2014.10.026 (2015).

36. Garyfallidis, E. et al. Dipy, a library for the analysis of diffusion mri data. Front. Neuroinformatics 8, DOI: 10.3389/fninf.2014.00008 (2014).

37. Mairal, J., Bach, F., Ponce, J. & Sapiro, G. Online learning for matrix factorization and sparse coding. J. Mach. Learn. Res. 11, 19–60 (2010).

38. Cook, P. A. et al. Camino: Open-source diffusion-mri reconstruction and processing (2006).

39. Clarke, J. Interhemispheric functions in humans: relationships between anatomical measures of the corpus callosum, behavioral laterality effects, and cognitive profiles. Thesis, University of California, Los Angeles (1990).

40. Rosenthal, R., Rosnow, R. L. & Rubin, D. B. Contrasts and effect sizes in behavioral research: A correlational approach. Contrasts and effect sizes in behavioral research: A correlational approach. (Cambridge University Press, New York, NY, US, 2000).

41. R Core Team. R: A Language and Environment for Statistical Computing. R Foundation for Statistical Computing, Vienna, Austria (2019).

42. Raffelt, D. A. et al. Connectivity-based fixel enhancement: Whole-brain statistical analysis of diffusion mri measures in the presence of crossing fibres. Neuroimage 117, 40–55, DOI: 10.1016/j.neuroimage.2015.05.039 (2015).

43. Raffelt, D. et al. Apparent fibre density: a novel measure for the analysis of diffusion-weighted magnetic resonance images. Neuroimage 59, 3976–94, DOI: 10.1016/j.neuroimage.2011.10.045 (2012).

44. Yeh, F. C. et al. Automatic removal of false connections in diffusion mri tractography using topology-informed pruning (tip). Neurotherapeutics 16, 52–58, DOI: 10.1007/s13311-018-0663-y (2019).

45. D’Souza, S., Ormond, D. R., Costabile, J. & Thompson, J. A. Fiber-tract localized diffusion coefficients highlight patterns of white matter disruption induced by proximity to glioma. PLoS One 14, e0225323, DOI: 10.1371/journal.pone.0225323 (2019).

46. Anderson, D. N., Duffley, G., Vorwerk, J., Dorval, A. D. & Butson, C. R. Anodic stimulation misunderstood: preferential activation of fiber orientations with anodic waveforms in deep brain stimulation. J Neural Eng 16, 016026, DOI: 10.1088/1741-2552/aae590 (2019).

47. Grussu, F. et al. Neurite dispersion: a new marker of multiple sclerosis spinal cord pathology? Ann Clin Transl Neurol 4, 663–679, DOI: 10.1002/acn3.445 (2017).

48. Sepehrband, F. et al. Brain tissue compartment density estimated using diffusion-weighted mri yields tissue parameters consistent with histology. Hum Brain Mapp 36, 3687–702, DOI: 10.1002/hbm.22872 (2015).

49. Harms, R. L., Fritz, F. J., Tobisch, A., Goebel, R. & Roebroeck, A. Robust and fast nonlinear optimization of diffusion mri microstructure models. Neuroimage 155, 82–96, DOI: 10.1016/j.neuroimage.2017.04.064 (2017).

50. Lebel, C. & Deoni, S. The development of brain white matter microstructure. Neuroimage 182, 207–218, DOI: 10.1016/j.neuroimage.2017.12.097 (2018).

51. Reynolds, J. E., Grohs, M. N., Dewey, D. & Lebel, C. Global and regional white matter development in early childhood. Neuroimage 196, 49–58, DOI: 10.1016/j.neuroimage.2019.04.004 (2019).

52. Jones, D. K., Knosche, T. R. & Turner, R. White matter integrity, fiber count, and other fallacies: the do’s and don’ts of diffusion mri. Neuroimage 73, 239–54, DOI: 10.1016/j.neuroimage.2012.06.081 (2013).

53. Kumar, R., Nguyen, H. D., Macey, P. M., Woo, M. A. & Harper, R. M. Regional brain axial and radial diffusivity changes during development. J Neurosci Res 90, 346–55, DOI: 10.1002/jnr.22757 (2012).

54. Qiu, D., Tan, L. H., Zhou, K. & Khong, P. L. Diffusion tensor imaging of normal white matter maturation from late childhood to young adulthood: voxel-wise evaluation of mean diffusivity, fractional anisotropy, radial and axial diffusivities, and correlation with reading development. Neuroimage 41, 223–32, DOI: 10.1016/j.neuroimage.2008.02.023 (2008).

55. Winston, G. P. et al. Advanced diffusion imaging sequences could aid assessing patients with focal cortical dysplasia and epilepsy. Epilepsy Res 108, 336–9, DOI: 10.1016/j.eplepsyres.2013.11.004 (2014).

